# Mapping single-cell transcriptomes to copy number evolutionary trees

**DOI:** 10.1101/2021.11.04.467244

**Authors:** Pedro F. Ferreira, Jack Kuipers, Niko Beerenwinkel

## Abstract

Cancer arises and evolves by the accumulation of somatic mutations that provide a selective advantage. The interplay of mutations and their functional consequences shape the evolutionary dynamics of tumors and contribute to different clinical outcomes. In the absence of scalable methods to jointly assay genomic and transcriptomic profiles of the same individual cell, the two data modalities are usually measured separately and need to be integrated computationally. Here, we introduce SCATrEx, a statistical model to map single-cell gene expression data onto the evolutionary history of copy number alterations of the tumor. SCATrEx jointly assigns cancer cells assayed with scRNA-seq to copy number profiles arranged in a copy number aberration tree and augments the tree with clone-specific clusters. Our simulations show that SCATrEx improves over both state-of-the-art unsupervised clustering methods and cell-to-clone assignment methods. In an application to real data, we observe that SCATrEx finds inter-clone and intra-clone gene expression heterogeneity not detectable using other integration methods. SCATrEx will allow for a better understanding of tumor evolution by jointly analysing the genomic and transcriptomic changes that drive it.

## 1 Introduction

Cancer is an evolutionary process in which cells accumulate mutations and form different subclones with potentially varying phenotypes [1, 2]. The diversity of cell states that make up a tumor is caused by genomic and epigenomic changes, as well as interactions with its microenvironment [3]. The resulting intra-tumor heterogeneity is a major cause of treatment failure and relapse [4, 5]. In particular, the manner in which different genomic changes, as opposed to other factors, contribute to cell-specific states is highly relevant in treatment decision making, but remains an open question [6, 2].

In recent years, a host of technologies to assay the genome, epigenome, transcriptome, and proteome of single cells have been developed [7]. These methods hold great potential for cancer research, as obtaining a multi-dimensional view of individual cancer cells is crucial to design targeted therapies [8]. While a few technologies allow for the joint measurement of two or more different types of molecular quantities, they are usually of much lower throughput and provide lower quality data in comparison to the respective single-modality protocols. This is the case for G&T-seq [9] and DR-seq [10] which, while simultaneously enabling copy number calling and gene expression quantification from the same single cell, are not scalable to beyond a few dozen cells. In contrast, droplet-based single-cell RNA-sequencing (scRNA-seq) [11] and Direct Library Preparation (DLP) [12] provide scalable transcriptomic and whole-genome sequencing assays, respectively, by trading off sequencing depth with cell throughput.

Without reliable and scalable experimental platforms to obtain joint RNA and copy number aberration (CNA) profiles from the same single cells, computational approaches can be used to match cells derived from the same sample but sequenced on separate platforms. Because single-cell CNA profiles can be grouped into subclones, the most natural approach is to assign single-cell transcriptomes to CNA subclones by assuming that gene expression is proportional to copy number. This approach is implemented in the tools Clonealign [13] and CCNMF [14]. However, these methods ignore the underlying evolutionary history of cancer cells, which may accumulate (epi)genomic aberrations as they divide. This hierarchical structure can be inferred by a number of computational methods [15], including for single-cell CNAs [16, 17]. Furthermore, cells within the same CNA subclone may have different expression profiles and form different groups of cell states as a result of other somatic mutations, epigenetic events or spatial effects [18]. While totally unsupervised clustering [19] and trajectory inference [20] methods for scRNA-seq may find spurious structure that does not respect the underlying subclonal structure, Clonealign and CCNMF are over-restrictive in forcing the clustering structure to be driven only by CNA subclones. The same limitation holds for Cardelino [21], an approach aimed at matching SNV subclones with scRNA-seq data.

Here, we present SCATrEx (single-cell augmentation of CNA trees with gene expression), a Bayesian model designed to find a hierarchical clustering structure in scRNA-seq data on top of a CNA tree. SCATrEx uses a nested tree structure based on the tree-structured stick breaking process [22] to augment a given CNA tree with nodes corresponding to non-CNA events, such as undetected mutations or epigenetic changes with consequences at the gene expression level. This enables a joint analysis of CNAs and gene expression at single-cell level (Figure 1).

**Figure 1:**
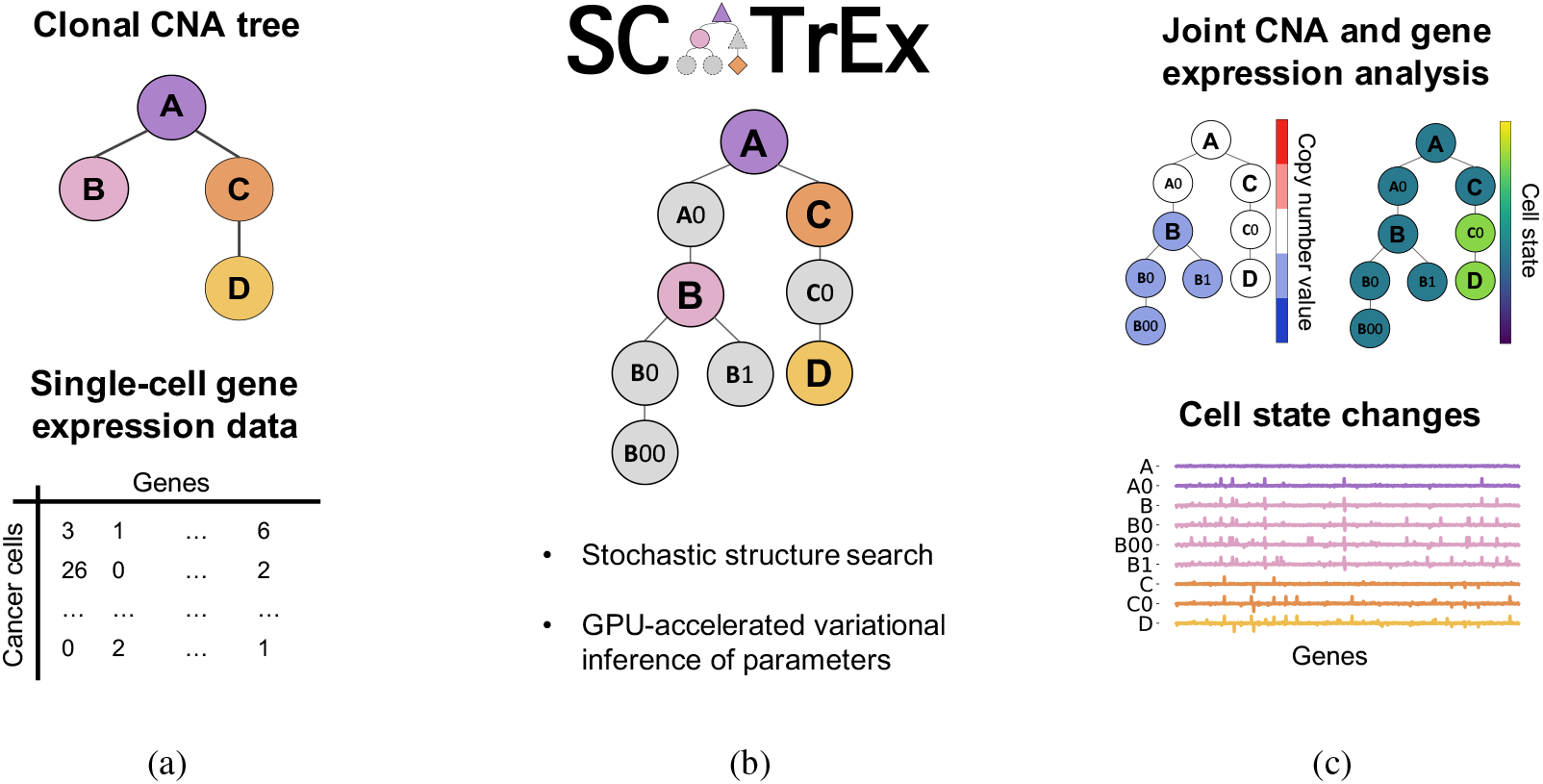
SCATrEx overview. (a) SCATrEx takes as input a copy number aberration (CNA) tree and single-cell gene expression data from cancer cells in matched tissue and produces (b) an augmented tree that contains CNAs and cell state changes. This enables an integrated view of genomic and transcriptomic heterogeneity in a tumor, in which gene expression clusters are guided by the known CNA clones. (c) SCATrEx further provides estimates of expression changes, including those not directly driven by the observed copy number values.

In Section 2, we introduce the model in detail, and in Section 3, we describe the inference procedure. In Section 4, we first show simulation studies indicating how the joint subclone assignment and clustering procedure of SCATrEx lead to robust assignments and improved clusterings under different levels of noise, in comparison to the fully-informed and the unsupervised approaches. We then apply SCATrEx to real breast cancer mouse xenograft data.

## 2 Model

SCATrEx models the gene expression heterogeneity of a tumor as a tree containing copy number events and transcriptional changes (Figure 1). To do this, we rely on three main assumptions. First, the CNA tree is fixed and known, and gene expression clusters live within each node of that tree. Second, these clusters are driven by CNAs and sparse changes in gene expression. Finally, these changes in gene expression may be caused by heritable changes of cell state, like SNVs or epigenetic events.

Formally, SCATrEx models each node in a known CNA tree as a tree-structured stick breaking process (TSSBP) [22]. We start this section by recalling the TSSBP, and then introduce our contributions: the nested tree-structured stick breaking process, and the observation model tailored to the scRNA-seq and CNA tree integration problem.

### 2.1 Tree-structured stick breaking process

The TSSBP defines a distribution over trees, where each node *∊* is associated with a mass *π_∊_* such that ∑_*∊*_*π*_*∊*_ = 1. These weights are constructed by interleaving two stick breaking processes defined by *ν_∊_*, which determines the size of the partition of node *∊*, and *ψ_∊_*, which determines the branching probabilities. Each node is labeled by a sequence of non-negative integers *∊* = *∊*_0_ *∊*_1_ …, with the root node being denoted by 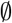. The children of the root are denoted by 0, 1, 2, … and the children of node 0 as 00, 01, 02, …, etc. The depth of node *∊* is the length of its sequence, *d_∊_* = |*∊*|. We define the mass assigned to each node by

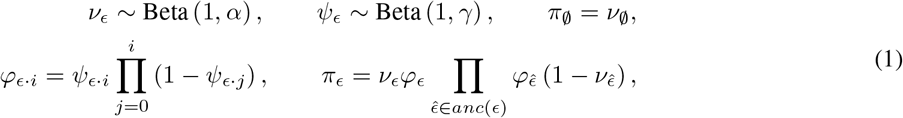

where 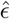 denotes the parent of *∊* and *anc*(*∊*) denotes the ancestors of *∊*. For two finite sequences *∊* and *∊*′, we denote by *∊* η *∊*′ their concatenation 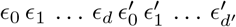, such that *∊* η *i* is the *i*-th child of *∊*, for all 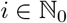.

The parameters *α* and *γ* control the shape of the tree: increasing *α* leads to deeper trees, and increasing *γ* increases the number of branches. In addition, if we wish to bound the depth or width of the obtained tree to specific values, we can truncate the *ν* and *ψ* sticks accordingly, yielding a truncated TSSBP. This is useful to define a distribution over weights of a tree with known, fixed structure.

### 2.2 Tree augmentation via nested tree-structured stick breaking

We now extend the TSSBP to model subclone-specific structure. Our approach relies on regarding the observed tree as a TSSBP in which each node is itself a tree defined by a TSSBP, whose structure is not known *a priori*.

Let *T* be a given tree indexed by nodes *τ*, where each *τ* is a letter in the English alphabet (Figure 2a). Here, each *τ* is itself a tree generated according to its own TSSBP (Figure 2b). We index the nodes of each *τ* by *τ* · *∊*, i.e., the sequence obtained by appending the index *∊* to the index *τ*. The nested tree-structured stick breaking process (NTSSBP) defines the mixture weights of *τ* · *∊* by defining a truncated TSSBP on *T* and another TSSBP for each *τ*. The weights of each *τ* in *T* are given by Equation (1) where we replace *∊* by *τ*, and we set *ν_τ_* = 1 if *τ* is a leaf node and *ψ_τ_* = 1 if *τ* does not have any subsequent siblings, as per the truncated TSSBP construction.

**Figure 2:**
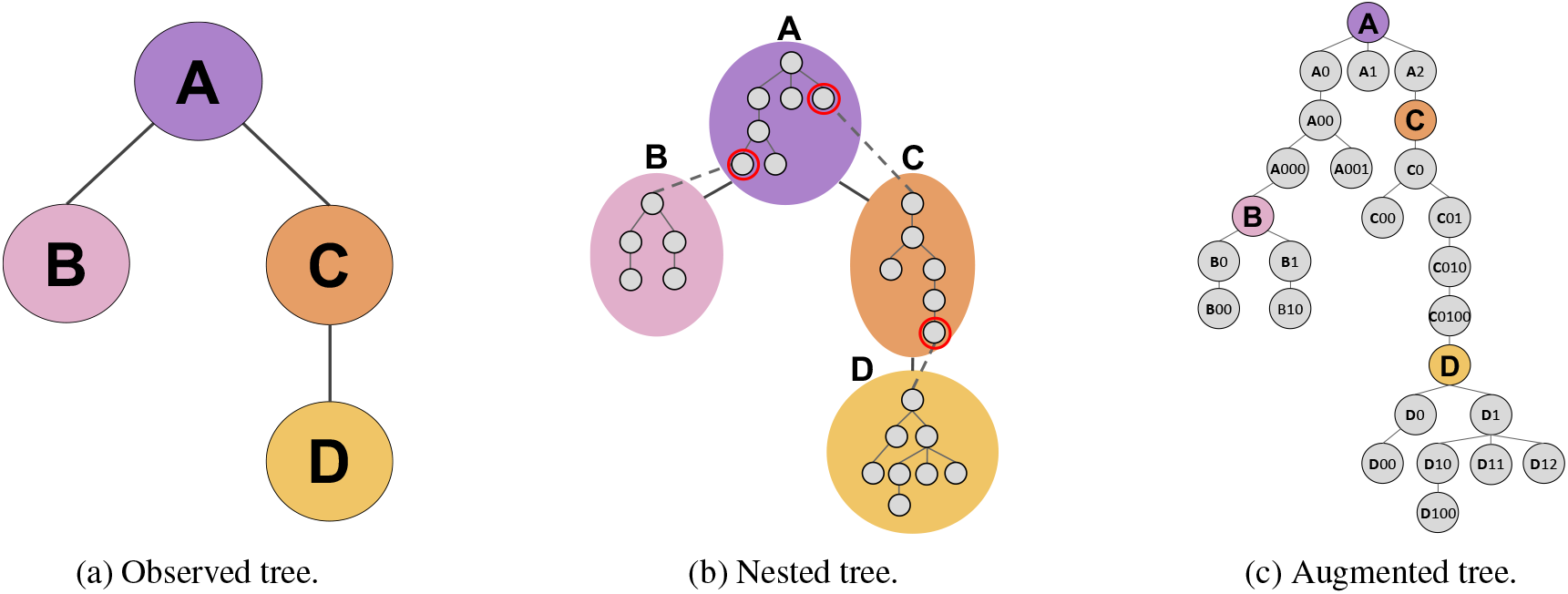
The tree augmentation model. (a) Coloured nodes denote known nodes in an observed tree (A, B, C and D). (b) The nested tree structured stick breaking process (TSSBP) generates a tree within each one of those nodes (gray circles and edges) through independent TSSBPs. A full tree is obtained by connecting (dashed gray edges) the root of each tree to a node (the pivot, highlighted in red) in its parent tree. (c) With the pivot connections in place, we obtain a full augmented tree, with the coloured nodes indicating the root of each TSSBP.

The weights of *τ* · *∊* conditioned on *τ* follow a *τ*-specific TSSBP, according to the following stick lengths:

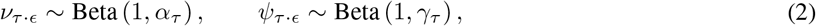

with weights defined as in Equation (1). These node weights sum to 1 conditioned on *τ* and we denote them by 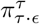. Then, the the weight of each node *τ* · *∊* in the full structure is given by

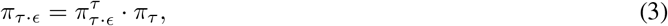

and we denote the process of obtaining the weights *π_τ·∊_* for all *τ* · *∊* as *π* ~ NTSSB (*ν, ψ, T*).

Finally, to obtain an augmented tree structure *T^a^* from the observed tree *T* and each of the *τ* trees, we connect the root 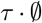 of each non-root *τ* to a node in its parent tree, 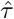. We call this connector node the pivot of *τ* and denote it *ρ_τ_*. While the independent TSSBP construction prevents the flow of mass from *ρ_τ_* to 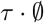, this connection serves to parameterize the observation model of 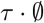 and obtain a single augmented tree structure. In general, the parent of node 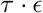 with 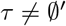 is 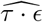, which corresponds to *ρ_τ_* if 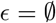 and 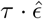 otherwise. For each non-root *τ*, its pivot is chosen according to a probability distribution that depends on the depth of each node in 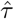 and a parameter *η_τ_*,

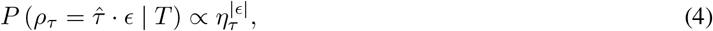

where *η_τ_* > 1 will put more mass in deeper nodes and *η_τ_* = 1 yields the uninformative prior. We denote the full tree obtained in this manner by the tuple (*T, ρ*) (Figure 2c). The depth of each node *τ* · *∊* in this tree is defined recursively as

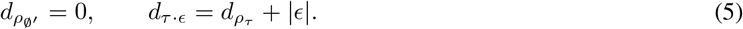

The nested tree construction provides a framework to encode diverse assumptions regarding the subclone proportions, the structure within each subclone, and the complexity between subclones.

The known subclone proportions can be plugged into Equation (3) via *π_τ_* directly. Otherwise, the parameters *α_T_* and *γ_T_* can be chosen so as to prefer some distribution of masses over others. In this case, we can also replace *α_T_* with a depth-dependent function like 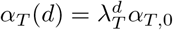, introduced in [22], where *d* denotes depth. This is useful to capture realistic cases such as when the root node in a CNA tree corresponds to diploid cells, which may not be present in late-stage tumours, and so we would like to assign a very low mass to it. The previous form for *α_T_*(*d*) can achieve this by setting *α*_*T*,0_ > 1 and λ ∈ (0,1).

The ability for each subclone to have their own expected structures is enabled by the usage of a separate TSSBP in each one, for which different hyperparameters (*α_τ_*, *γ_τ_*) can be set. Finally, the complexity between subclones is controlled by the pivot nodes, and each subclone tree *τ* can prefer to inherit the data-emission parameters from more or less deep nodes in its parent 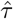, according to the hyperparameter *η_τ_*. For example, it is likely that large genomic rearrangements between subsequent nodes in a CNA tree are driven by some mechanism of genomic instability that could be detected from gene expression [23]. Thus, if a CNA tree has a node with a large number of events, it may be appropriate to bias the augmentation model towards adding nodes between that node and its parent.

### 2.3 Observation model to augment CNA trees with scRNA-seq data

We now turn to define the data-generating process of each node *τ* · *∊* in each tree *τ* in *T^a^*. In general, we assume each *τ* is equipped with a known data-generating parameter *x_τ_*. The unobserved nodes *τ* · *∊* generate data according to a probability distribution with mean dependent on *x_τ_* and unobserved node-specific parameters.

We wish to model single-cell RNA-seq data obtained from the same tumor as the one from which a CNA tree *T* was derived. Each observed node *τ* is a subclone for which a copy number profile x_*τ*_ for all genes is known. The data is collected in a cell-by-gene matrix **Y**. For each cell *n* = 1, …, *N* and each gene *g* = 1, …, *G*, observation *y_ng_* is the number of reads from cell *n* that mapped to gene *g*, with total read counts per cell denoted by *s_n_*. The observations are generated according to the following process:

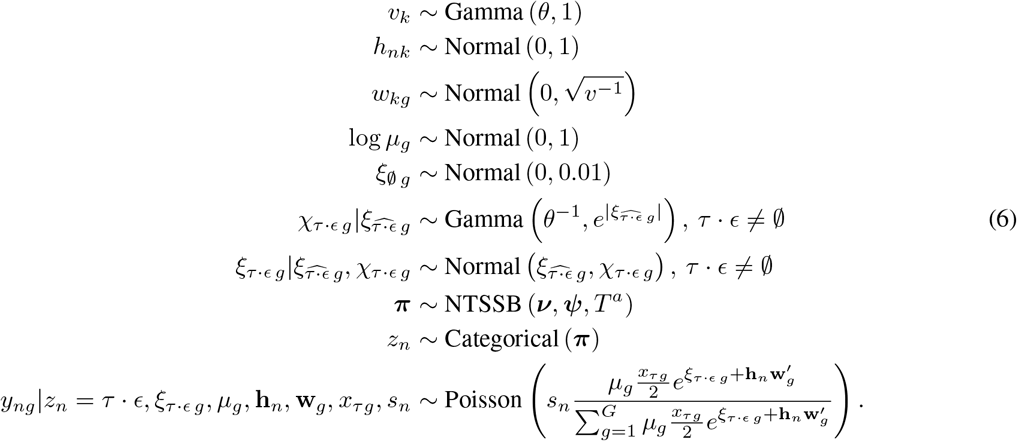

In this model, the gene expression of a cell is proportional the copy number profile *x_τ_* of the subclone it belongs to and influenced by node-specific cell state factors *ξ*_*τ*·*∊*_ that are propagated through the augmented tree. Additional structured noise factors **hw**′ contribute to gene expression variability that does not follow the tree structure.

The unobserved node-specific data-generating parameters are *ξ* and **χ**. The variable *ξ*_*τ·∊ g*_ captures gene expression changes along the evolutionary history of the tumor which are not captured by the observed CNA tree. The Normal-Gamma mixture prior on non-root nodes *ξ*_*τ·∊ g*_ is built to capture two properties: sparsity and hard inheritance of the previous events. Sparsity enforces that each unobserved event affects only a small number of genes. Inheritance is enforced by centering the expected value at the previous event and favouring the appearance of new events in previously unaffected genes. We have omitted the dependence of these terms on the pivot node assignments ***ρ*** as it is clear that if *τ* · *∊* is such that 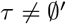 and 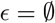, then 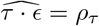.

In addition, ***h*** and ***w*** denote a matrix decomposition of the data into *K* factors, which is used to separate confounding factors from the tree-dependent terms. *μ_g_* is a gene scale factor indicating the per-copy expression of gene *g*, which allows us to relate the expression level with τ’s copy number state by multiplying *μ_g_* by 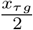, where *x_τg_* = 2 indicates that subclone *τ* is diploid for gene *g*.

We additionally couple the prior distributions on the scales of the noise factors *v_k_* and the shape of ***χ*** in the Normal-Gamma mixture via a hyperparameter *θ*. Specifically, we want to ensure that not only we maintain sparsity on *ξ*, but also that large events have a higher density under *ξ* than under the noise factors. By default, we set *θ* to 100.

The model (6) can be seen as a generalization of Clonealign [13], which defines a simple mixture model with number of components equal to the number of observed clones, with additional structured noise. In other words, our model reduces to Clonealign by removing the node-specific variables that define cell state events (***ξ*** and ***χ***) and drawing the assignments *z_n_* from a Categorical distribution over the known clones with no prior distribution on the mixture components. As in [13], we normalize the mean expected read counts by total RNA to model the relative abundances of each gene within a cell and scale by *s_n_*.

## 3 Inference

Given data **Y** and an observed tree *T*, we are interested in learning the augmented tree structure and its parameters. We use a search-and-score approach to structure learning and approximate the score of each augmented tree with a mean-field variational method. Formally, we are interested in finding the augmented tree (*T^a^*, ***ρ***) such that the joint probability of data and ***ρ*** given *T^a^* is maximized:

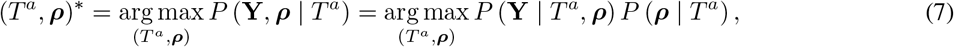

where *P*(**Y**|*T^a^*, ***ρ***) is the marginal likelihood of the data and *P*(*ρ*|*T^a^*) is the prior probability distribution on the pivot node assignments. For the optimal tree (*T^*a*^*, ***ρ***)*, we estimate the values of the latent variables as their posterior means given **Y**. In the following sections we describe the details of this optimization.

### 3.1 Marginal likelihood approximation

In a non-conjugate model with latent variables such as the one in Equation (6), finding the marginal likelihood of data for a given structure is analytically intractable. The main approaches to obtain this quantity computationally are sampling methods such as Markov chain Monte Carlo (MCMC) [24] and variational inference [25]. Because we are interested in finding the best *T^a^* that maximizes the marginal likelihood, we need to run this approximation many times, once for each tree. We use variational methods, because these are often able to obtain an approximation faster than MCMC [26, 27]. This is the approach used in [28], for example.

Here, we use a mean-field automatic differentiation variational inference (ADVI) approach as introduced in [29]. Variational inference maximizes a lower bound on the marginal data likelihood (the so-called evidence lower bound, i.e., the ELBO) by approximating the posterior distribution over the latent variables in the model. The posterior is approximated with a parametric family of distributions whose parameters are optimized in order to maximize the ELBO.

In ADVI, this optimization is gradient-based, so we write the likelihood in terms of continuous variables only, by marginalizing out the discrete variables *z_n_* in Equation (6). We thus weigh the likelihood of data given node *τ* · *∊* by its mixture weight *π*_*τ*·*∊*_ and sum over all nodes:

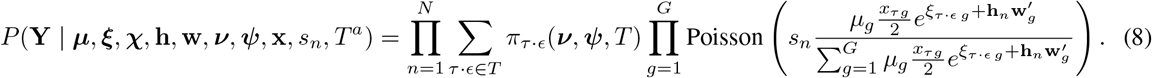

We approximate the posterior distribution over all the latent variables in the model with a mean-field (i.e., fully factorized) family of variational distributions:

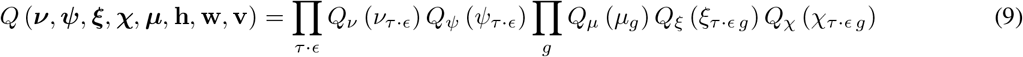

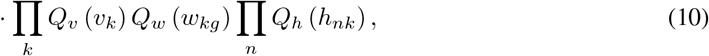

and use Normal distributions for all factors in unconstrained space. We constrain *ν* and *ψ* to the [0,1] range via the softmax function, and *χ* and *υ* to the positive reals via the exponential function. The variational parameters are then the means and standard deviations of these Normal distributions.

With the variational distribution in place, we can summarize the optimization problem for each tree as:

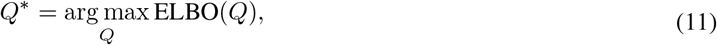

with

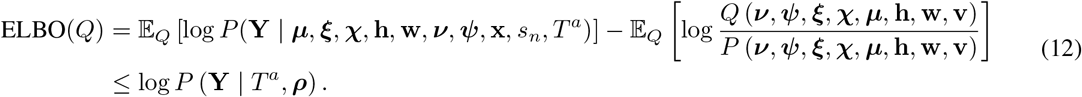

The use of the mean-field approximation comes with a disadvantage, as this objective function does not penalize copies of nodes. The model states that each node’s *χ* puts most density near zero, and its *ξ* is centered at the parent’s value. The maximum likelihood solution is then to set each node to copy exactly the parent’s with *χ* equal to zero, which would lead to having lots of nodes with equal *ξ*. In principle, this would be avoided by the second term in Equation (12), by penalizing low entropy variational distributions such as the ones resulting from such a degenerate case. By ignoring the posterior correlation between the different nodes, the mean-field approximation is unable to apply this penalty. Thus, in this paper we simply penalize those copies by adding a term to Equation (12) for each node that is equal to the prior probability of that node’s *ξ* being exactly equal to the parent, and *χ* being very close to zero in all genes.

The non-conjugate nature of the model in Equation (6) does not allow us to compute the ELBO analytically, so following [29] we draw Monte Carlo samples from *Q* as an approximation, and employ reparameterization gradients to update the variational parameters in a stochastic gradient ascent manner. We implement this optimization scheme using the automatic differentiation Python library JAX [30], which enables parallelization on GPU to achieve very fast runtimes, and use the widely adopted Adam optimizer [31].

### 3.2 Structure learning

We move in the space of trees by proposing changes to the tree, optimizing the ELBO for the new tree and accepting the changes if the total score (i.e., weighing the ELBO by the pivot prior as in Equation (7)) improves. We use 5 discrete moves for structure search. The basic add and merge moves are used to expand and contract the model size (as in [28]) whereas the remaining moves rearrange nodes and subclone trees. To move more efficiently, some moves are more likely than others, and their probability may depend on the current state of the tree, such as the proportion of data explained by each node. We additionally interleave the structure updates with updates to the baseline and global noise factor parameters, **λ**^*μ*^, **λ**^*w*^ and **λ**^*h*^, with all the node-specific variational parameters fixed.

At each step, we only optimize the ELBO with respect to the variational parameters of the nodes that were affected by the proposed move. We run the ELBO optimization for a proposed tree until a maximum of 500 steps or until the change in the average ELBO across a window of size 50 is less than 10^−6^. We detail the moves in the following paragraphs.

#### 3.2.1 Add

This move chooses a node *τ* · *∊* anywhere in the tree and creates a new child *τ* · *∊* · *i*. The probability of choosing *τ* · *∊* is proportional to the number of data points best explained by it. When we add *τ* · *∊* · *i*, we collect the data most explained by its parent *τ* · *∊* in 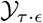. If *τ* · *∊* has no other children, we initialize *ξ*_*τ*·*∊·i*_ such that it is centered at the datum in 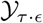 which allocates the smallest likelihood on *τ* · *∊*. Then we update the variational parameters of *τ* · *∊* and *τ* · *∊* · *i* until convergence.

We accept the new child irrespective of the change in ELBO, and rely on subsequent merge moves to remove redundant nodes. This provides an effective method to escape local optima. However, if the number of nodes with mass *π*_*τ*·*∊*_ smaller than 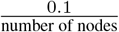 is greater than 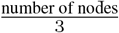, we assume there are probably too many nodes in the tree and we turn to only accepting a birth if the ELBO improves.

#### 3.2.2 Merge

The merge move first selects tree *τ*, weighed by the number of nodes it contains, and then chooses a pair of nodes within *τ* to merge. The pair is chosen by first selecting a non-root node *τ* · *∊_a_*, and then assessing which one of its siblings and parent contains the parameter *ξ*_*τ*·*∊_b_*_ at the lowest mean-squared distance from *ξ*_*τ*·*∊_a_*_. Upon merge, the children of *τ* · *∊_a_* are moved to *τ* · *∊_b_*, and the variational parameters of the two nodes are combined into *τ* · *∊_b_* as

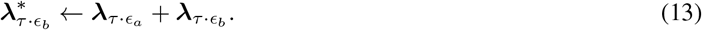

We then iterate the variational parameters of all nodes below *τ* · *∊_b_* until convergence and accept the move if the ELBO improves. This move removes redundant nodes and relies on the stick length priors to penalize nodes which do not add sufficient explanatory value to the tree to compensate for the added tree complexity.

#### 3.2.3 Swap

In this move, we select two nodes from the augmented tree *τ* · *∊_a_* and *τ* · *∊_b_* and swap their variational parameters, i.e.,

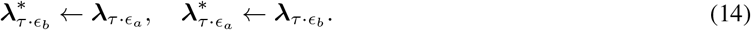

When swapping 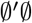 with one of its children 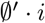: the root node’s mean is dominated by the baseline expression, so we update the baseline and all the unobserved factors in the tree simultaneously until convergence of the ELBO. This leads to a large change in the variational parameters, and takes a longer time to converge than the other moves.

In addition, if one of the swapped nodes is the pivot of a subtree, we update the pivot assignment such that it remains in the same location as before. In other words, if 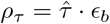 and 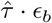 is swapped with 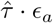, we set 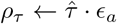.

#### 3.2.4 Pivot prune and reattach

This move uniformly chooses a non-root *τ* and proposes a pivot *ρ_τ_* from its parent, 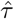. The probability of choosing 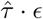 as the pivot is proportional to the inverse mean-squared distance between 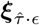 and *ξ*_*τ*·*∊*_, weighed by the pivot prior probability defined in Equation (4). We update the variational parameters of all the nodes in *τ* and accept the change if the total score improves.

#### 3.2.5 Subtree prune and reattach

This move chooses *τ_a_* and *τ_b_* uniformly from the observed nodes. From *τ_a_* it prunes some node *τ_a_* · *∊* and its children and reattaches that subtree to *τ_b_*. This move is used to escape local optima in which unobserved factor lineages are in the incorrect *τ*. This move can produce a dramatic change to the tree and is thus proposed at a lower frequency compared to the others.

#### 3.2.6 Intra-subtree prune and reattach

This move chooses one node *τ* · *∊_a_* from any subtree *τ* and reattaches it to a different parent *τ* · *∊_b_* than the one it currently has, within the same subtree. Node *τ* · *∊_b_* is chosen from the possible nodes in *τ* according to the similarity of their current variational means for *ξ*.

## 4 Results

### 4.1 Simulated data

We assess the ability of SCATrEx to find the ground truth clusters and clone assignments in a simulation setting where we vary the number of noise factors and the number of nodes per clone. Our procedure begins by simulating a copy number tree containing 4 nodes with copy number profiles defined on 30 genomic regions of unequal size. This tree is then augmented by adding 0, 1 or 2 extra nodes to each copy number clone corresponding to gene expression clusters. For this augmented tree structure, we generate 1000 gene expression profiles across 800 genes according to Equation (6), and we vary *K* between 0, 2 and 4.

To measure the ability of SCATrEx to recover all the simulated clusters, we use the Adjusted Rand Index [32] (Figure 3a). As there is currently no other method that uses the CNA tree to guide the inference of clusters from scRNA-seq data, we compare against two other simple alternatives. First, we simply apply Leiden [33] which is the current standard clustering method for scRNA-seq data. To imbue Leiden with prior knowledge about the CNA clones, we also report the performance of first running Clonealign [13] to identify the clones, and then running Leiden within each clone.

**Figure 3:**
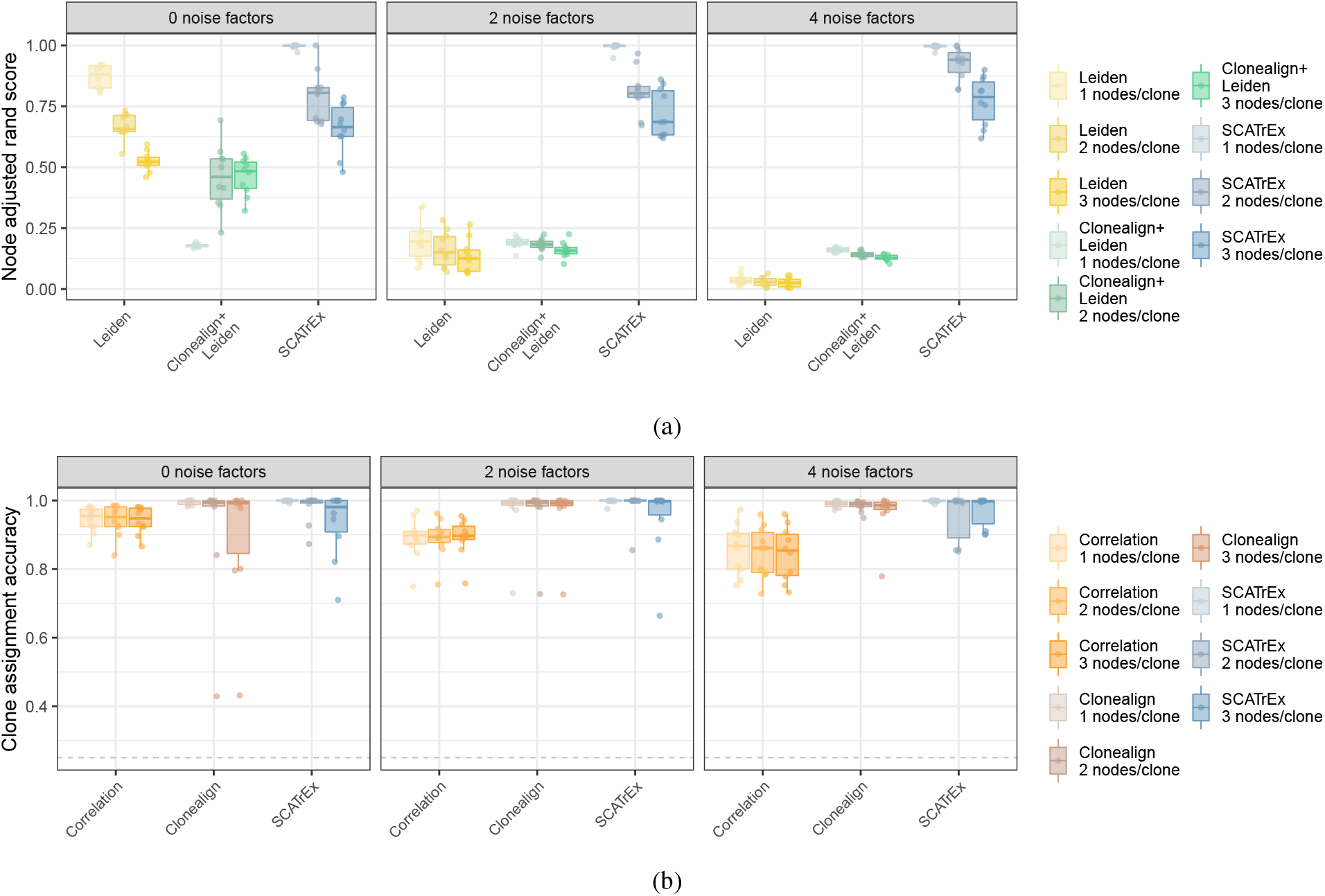
Performance assessment of clone assignments and clustering in simulated data. For random augmented trees with 4 copy number clones, and varying the number of nodes per clone from 1 to 3 (resulting in a total of 4, 8 and 12 total nodes), we generated noisy gene expression data for 1000 cells and 800 genes. The number of noise factors was set to 0, 2 and 4. (a) We compared the clustering performance of SCATrEx (blue) with the totally unsupervised method Leiden (yellow), and an *ad hoc* clone-specifc clustering approach by applying Leiden within each clone assigned by Clonealign (green), which does not assume any tree structure is present. (b) We additionally compared the accuracy of the cell to clone assignments with a naive correlation-based method (orange) and Clonealign (brown). We also show the random assignments as baseline (gray dashed line).

Leiden performs well when *K* = 0 but quickly deteriorates as its model-free approach is unable to distinguish the signal that forms the different clusters from the structured noise. Restricting it to the clone assignments of Clonealign (Clonealign+Leiden) gives a slight improvement in the noisy settings, but this two-step approach is still far from achieving the performance of SCATrEx, which draws strength from the joint modelling of noise, clones and clusters.

In addition to accurately reconstructing the true clustering structure, SCATrEx matches the performance of Clonealign in the task of assigning cells to the correct CNA clones of origin, as measured in terms of assignment accuracy (Figure 3b). We compare SCATrEx with Clonealign, which can be thought of as a reduced version of SCATrEx – i.e., without the NTSSB prior and the cell state factors. As a baseline, we report the accuracy achieved by assigning each cell to the CNA profile that its normalized gene expression most strongly correlates with.

As expected, the results show that as either the number of noise factors or the number of nodes per clone increase, the worse the naive correlation method performs. In contrast, both Clonealign and SCATrEx are robust to the noise level, with the number of nodes per clone being the main source of variance in performance.

We find that SCATrEx is overall better than unsupervised clustering approaches while matching the state-of-the-art clone assignment performance. The sub-optimal performances reported in Figure 3 are due to cases in which, even though each data set was simulated to contain a fixed number of nodes and each node forced to have cell state factors, their scales (as generated from the model in Equation (6)) were too small to be detectable. We also observed that when there are very large cell state events overlap with CNA events near the root of the tree, the assignments can be confounded.

For each simulated data set, we ran both Clonealign and SCATrEx with 5 different initializations and for 500 steps each, and report the run that achieved the highest ELBO. We used the default parameters for Leiden as provided by the scanpy package [34].

### 4.2 Real data

We applied SCATrEx to the data set introduced in [13, 12] of independent measurements of scRNA-seq and scDNA-seq from a mouse xenograft of a triple-negative breast cancer tumor. The authors reconstructed an evolutionary tree that relates three copy number subclones A, B and C, where A features a chromosome X deletion, and B and C share very similar copy number profiles.

The results are shown in Figure 4. SCATrEx inferred a tree containing various extra clusters within the observed CNA nodes, and identified cell state events in all nodes in the augmented tree (Figure 4a). We used the strategy applied in [13] to validate the assignments, which is to plot the mean log expression of each copy number state, and observe a positive correlation, as expected (Figure 4b, top).

**Figure 4:**
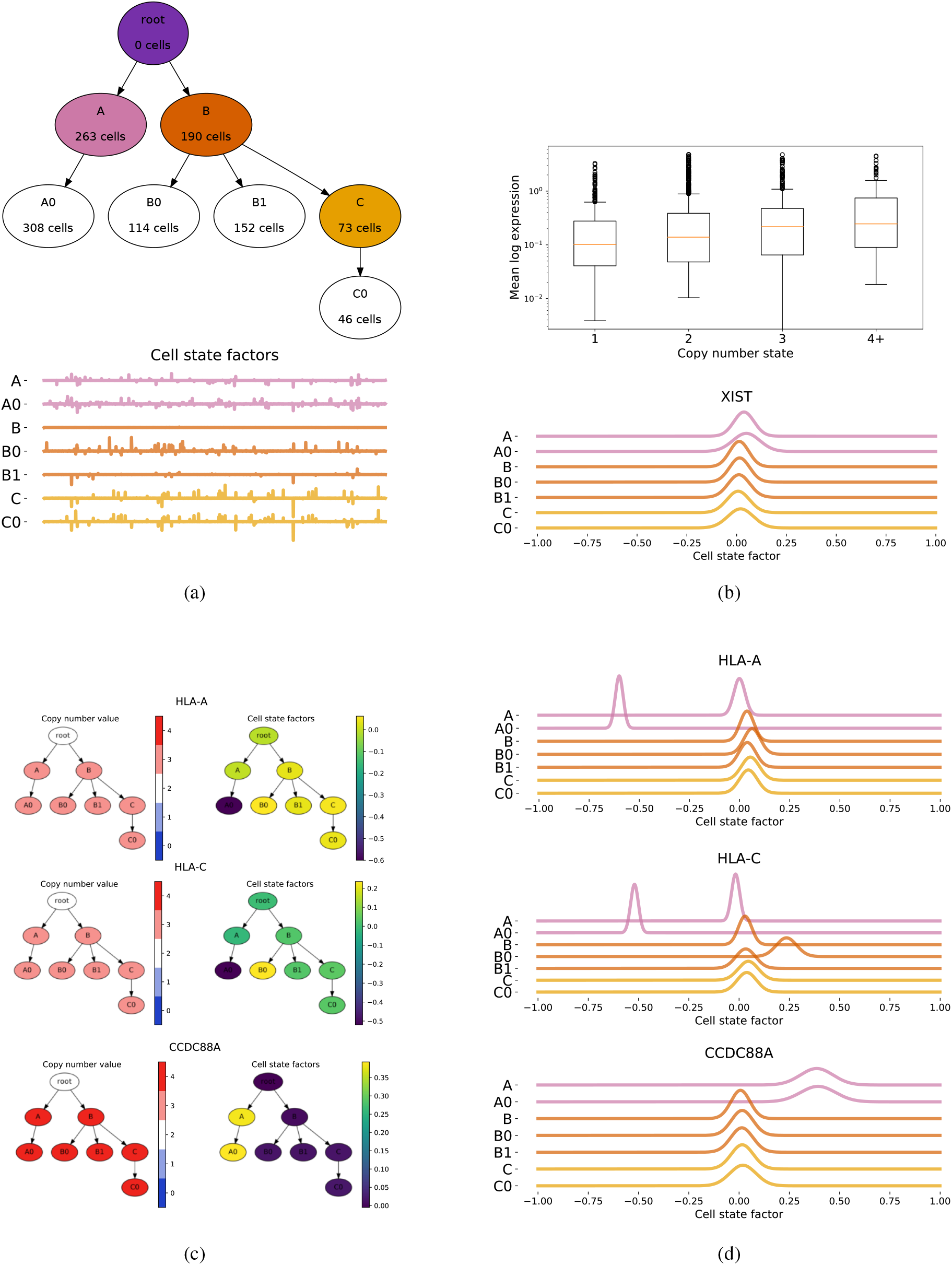
Application to data from a mouse xenograft of a triple negative breast cancer. (a, top) SCATrEx learned a tree with extra clusters within clone A, B and C, with (a, bottom) its nodes containing various cell state events. (b, top) Based on the SCATrEx cell-to-clone assignments, the correlation between the mean log expression and the copy number states is positive as expected. (b, bottom) The posterior distribution of the cell state factor for gene XIST shows its deletion is the cause of its underexpression. (c) Mapping the CNA and cell state factors of specific genes on the tree shows complementary signals contributing to gene expression. (d) The posterior distributions of the cell state factors for the same genes.

Additionally, we assessed whether the inactive chromosome X-specific gene, XIST, was affected by any cell state event. In Figure 4b (bottom) we plot the variational distribution of *ξ* for this gene in every node, and do not observe any significant change. Since the gene was observed to be lowly expressed in subclone A [13], this confirms that the lost copy was the inactive one. Otherwise, SCATrEx would reveal an up-regulation of the gene in this subclone.

Analysing the CNA and cell state events side by side for a few selected genes provides insights into possible mechanisms of tumor growth. We highlight three genes: HLA-A, HLA-C and CCDC88A. These genes have the same CNA across all subclones, but are affected differently by cell state changes. HLA-A and HLA-C are down-regulated in cluster A0 of clone A, which is partly consistent with the analysis of [13]. However, here we have the power to distinguish subcluster-specific expression, and find that this down-regulation is not shared across all cells in clone A. In contrast, CCDC88A was previously reported to be up-regulated in subclone A, and here we confirm this by observing that this is shared across all cells in subclone A. Like the authors of the original study, we can not tell whether these expression changes are *in trans* regulatory effects of CNAs, the product of other somatic mutations, or epigenetic changes. Nonetheless, SCATrEx readily provides these results as part of its inference, without requiring further differential expression testing, in contrast to the approach in [13].

## 5 Discussion

Resolving intra-tumor heterogeneity at single-cell level requires the integration of multiple molecular measurements. To make this integration interpretable, it is important to incorporate prior knowledge in the methods that tackle this task. Here we developed a novel framework to model single-cell gene expression in the context of a given clonal copy number tree, enabling us to jointly assign cells to their clones of origin and find clone-specific clusters. When learning this structure, the possible gene expression variability that does not arise from it needs to be accounted for. Our model therefore considers three types of variation: tree-structured known copy numbers, tree-structured unknown sparse gene expression changes and unknown noise factors that do not follow a tree structure.

Simulations demonstrate that our joint approach is accurate in both cell-to-clone assignments and clustering. In a real data example of a mouse xenograft of triple-negative breast cancer, we obtained an augmented tree and inferred gene expression states for the clones as well as for clone-specific populations of cells. Such expression states may be the result of unseen genomic or epigenomic changes, but our model does not have the ability to identify their root cause.

The cell populations present in tumors typically do not include the cell of origin that contains the healthy gene expression profile. This makes defining the baseline expression levels difficult. While this issue is attenuated by the use of prior distributions on the model parameters, it can still lead to local optima in the model inference. It is therefore crucial that different initialisations are tried for every SCATrEx run.

SCATrEx works with a large number of genes to extract the most information from the data, which leads to increased computational costs. However, retaining a smaller number of highly variable genes could result in removing genes that are responsible for smaller groups of cells. Additionally, the sparse nature of the cell state signals captured by our model does not necessarily mean they consist of highly variable genes across the whole data set.

While SCATrEx forces the clustering structure to respect the known CNA subclones, there may be cells with similar expression profiles that belong to different subclones. This can happen, for example, if spatial factors have a larger influence on the gene expression of cancer cells than their CNA profiles. Instead of being attributed nodes in the SCATrEx tree, these clusters may be captured by the structured noise factors.

Inferring trajectories of tumor development in cancer has been a longstanding goal in cancer research. While clonal tree reconstruction provides detailed genomic information, the inferred lineages do not reflect the full extent of phenotypic heterogeneity. On the other hand, trajectories inferred from single-cell transcriptomics do not necessarily reflect genomic relationships [35]. A combined representation of intra-tumor heterogeneity and its evolution, enabled by SCATrEx and future derivations, will help advance cancer biology.

## Code availability

The Python code for SCATrEx and Snakemake [36]workflows for the analysis shown in this manuscript are publicly available at https://github.com/cbg-ethz/SCATrEx.

## Acknowledgements

Part of this work was supported by SNSF Grant 310030_179518 (http://www.snf.ch) and by ERC Synergy Grant 609883 (http://erc.europa.eu/).

